# Evolution of cytokine production capacity in ancient and modern European populations

**DOI:** 10.1101/2021.01.14.426690

**Authors:** Jorge Domínguez-Andrés, Yunus Kuijpers, Olivier B. Bakker, Martin Jaeger, Cheng-Jian Xu, Jos W.M. van der Meer, Mattias Jakobsson, Jaume Bertranpetit, Leo A.B. Joosten, Yang Li, Mihai G. Netea

## Abstract

As our ancestors migrated throughout the different continents, natural selection increased the presence of alleles advantageous in the new environments. Heritable variations that alter the susceptibility to diseases vary with the historical period, the virulence of the infections, and their geographical spread. In this study we built polygenic scores for heritable traits influencing the genetic adaptation in the production of cytokines and immune-mediated disorders, including infectious, inflammatory, and autoimmune diseases, and applied them to the genomes of several ancient European populations. We observed that the advent of the Neolithic was a turning point for immune-mediated traits in Europeans, favoring those alleles linked with the development of tolerance against intracellular pathogens and promoting inflammatory responses against extracellular microbes. These evolutionary patterns are also associated with an increased presence of traits related to inflammatory and auto-immune diseases.

## Introduction

Human history has been shaped by infectious diseases. Human genes, especially host defense genes, have been constantly influenced by the pathogens encountered (Fumagalli and Sironi, 2014; Karlsson et al., 2014; Quintana-Murci and Clark, 2013). Pathogens drive the selection of genetic variants affecting resistance or tolerance to the infection, and heritable variations that increase survival to diseases with high morbidity and mortality will be naturally selected in people before reproductive age (Karlsson et al., 2014). These selection signatures vary with historical period, virulence of the pathogen, and the geographical spread.

Here we investigated the historical evolutionary patterns leading to genetic adaptation in cytokine production and immune-mediated diseases, including infectious, inflammatory, and autoimmune diseases. Cytokine production capacity is a key component of the host defense mechanisms: it induces inflammation, activates phagocytes to eliminate the pathogens and present antigens, and controls induction of T-helper adaptive immune responses. We have therefore chosen to investigate the evolutionary trajectories of cytokine production capacity in modern human populations during history. To determine the difference in polygenic regulation of diseases and cytokine production capacity, we used data derived from the 500 Functional Genomics (500FG) cohort of the Human Functional Genomics Project (HFGP; http://www.humanfunctionalgenomics.org). The HFGP is an international collaboration aiming to identify the host and environmental factors responsible for the variability of human immune responses in health and disease (Netea et al., 2016). Within the HFGP project, the 500FG study generated a large database of immunological, phenotypic and multi-omics data from a cohort of 534 individuals of Western-European ancestry, which have been used to integrate the impact of genetic and environmental factors on cytokine production and immune parameters. We subsequently deciphered the factors that influence inter-individual variation in the immune responses against different stimuli (Bakker et al., 2018; Li et al., 2016; Schirmer et al., 2016; Ter Horst et al., 2016).

## Results and Discussion

Peripheral blood mononuclear cells from these individuals were challenged with bacterial, fungal, viral and non-microbial stimuli, and six cytokines (TNFα, IL-1β, IL-6, IL-17, IL-22 and IFNγ) were measured at 24h or 7 days after stimulation, generating 105 cytokine-stimulation pairs (Fig. S1 and Table S1). We correlated cytokine production with genetic variant data to obtain cytokine quantitative trait loci (QTLs), which were employed to compute and compare the polygenic risk score (PRS) of the genomes of 827 individuals from different human historical eras (early upper Paleolithic, late upper Paleolithic, Mesolithic, Neolithic, post-Neolithic) which were downloaded from version 37.2 of the compiled dataset containing unimputed published ancient genotypes (https://reich.hms.harvard.edu/downloadable-genotypes-present-day-and-ancient-dna-data-compiled-published-papers), and 250 modern Europeans randomly selected from the European 1000G cohort (see accompanying manuscript by Kuijpers et al.). We then investigated how the PRS changes over time by constructing linear models and performing correlation analysis. In order to account for the ancient DNA samples being pseudo-haploid, ambiguous SNPs (A/T and C/G) were excluded when computing PRS to prevent errors due to strand flips. PRS was computed using the most significant QTLs that had a P value lower than our predetermined threshold for each given trait and removing all variants within a 250kb window around these variants. The dosage of these variants was multiplied by their effect size while the dosage of missing variants in a sample were supplemented with the average dosage. Finally, we scaled the PRS to a range of −1 and 1 and correlated the scores of the samples with their respective carbon dated age. In order to verify the robustness of our results we repeated the analysis at multiple threshold combinations for variant missingness and QTL thresholds. Furthermore, an analysis-based down-sampling approach shows that the trajectories observed in our results are consistent regardless of the sample size (Fig. S2). A schematic representation of the steps performed is shown in Fig. S3.

Applying the methodology described above, several patterns were apparent (Fig. 1). The first overall observation is that the estimation of cytokine production capacity based on PRS shows significant differences between populations in various historical periods, and the strength of evolutionary pressure on cytokine responses was different before and after the Neolithic revolution. We did not observe significant changes in cytokine production capacity between individuals who lived at different historical periods before the Neolithic, whereas strong pressure is apparent after adoption of agriculture and animal domestication in Europe. This different pattern may have resulted from the more limited number of samples available for the older time periods, resulting in lower statistical power, but the presence of some evolutionary pressure also before the Neolithic argues that this is most likely not the full explanation. The development of agriculture and domestication of animals in the Neolithic increased population densities on the one hand, and the contact between humans and domesticated animals as source of pathogens on the other hand. The number of zoonoses increases dramatically (examples being tuberculosis, brucellosis, Q-fever, and influenza), which strongly increased the selective pressure and caused significant adaptations of immunity at the genetic level (Flandroy et al., 2018). Most of the genetic adaptations to pathogens took place in the period since modern humans abandoned their hunting-gathering lifestyle and developed agriculture (Deschamps et al., 2016). In this respect, the strongest changes leading to tolerance (decreased cytokine production) were exerted in the cytokine responses to intracellular zoonotic infections (tuberculosis and *Coxiella*) (Fig. 1). In contrast, responses to the extracellular pathogens *Staphylococcus aureus* and *Candida albicans* indicate increased resistance, with high production of IL-22 and TNFα, respectively. The increased response to the important fungal pathogen *C. albicans* after the Neolithic period is validated also at transcriptional level. Overall, these patterns are reminiscent of the studies showing that human immune responses need to adapt to a new landscape of infectious agents depending on geographical location and types of microbe encountered (Ferwerda et al., 2007). Such different patterns were most likely encountered also through history.

**Figure 1:**
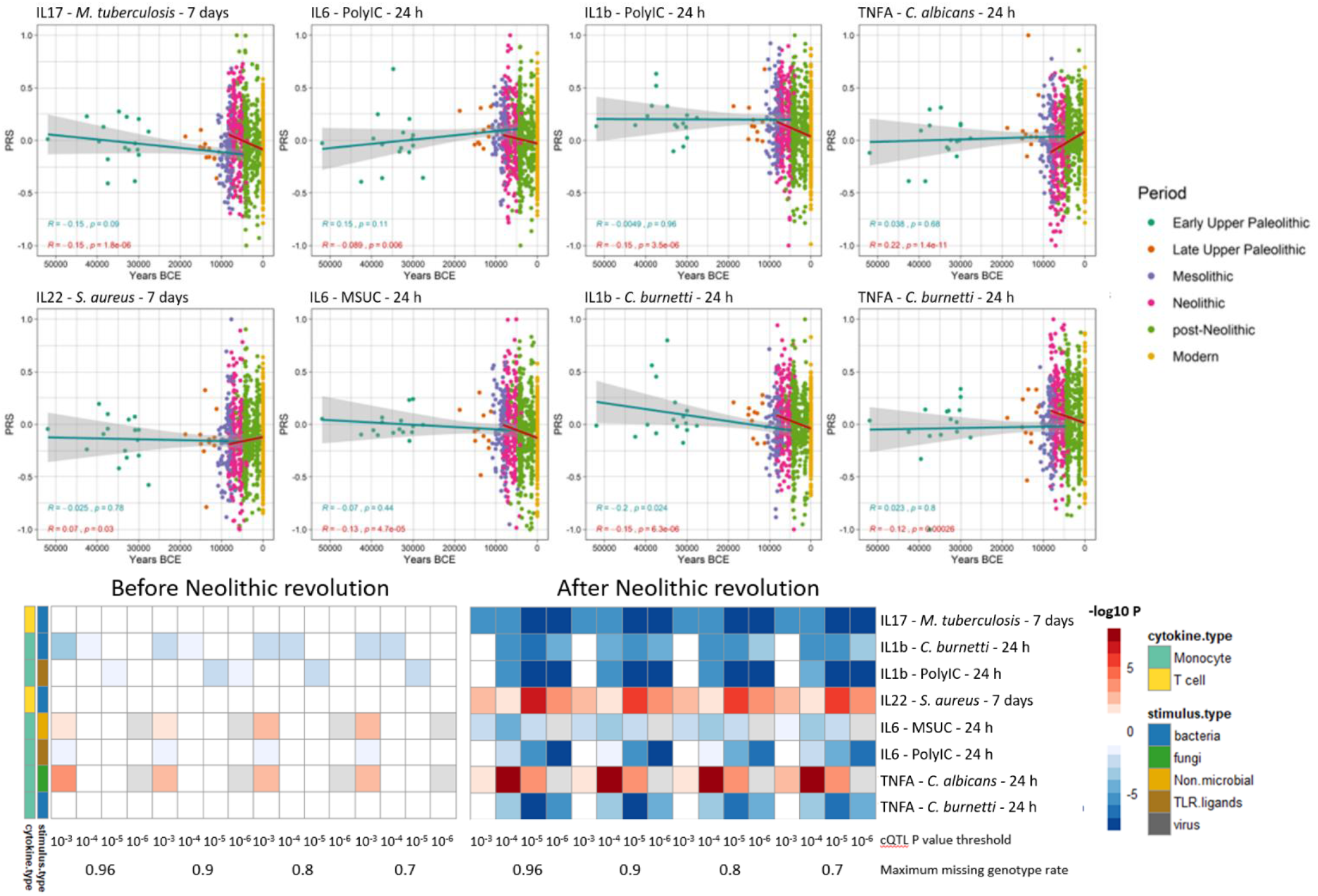
A) Correlation between cytokine PRS and time. Samples are colored by broad age period. The blue regression lines show PRS before the Neolithic revolution remained relatively constant for all traits whereas the red regression lines show the correlation after the start of the Neolithic period. The threshold of max missing genotype per sample was 0.96 and QTL P value cutoff was 10^−4^. MSUC: Monosodium urate crystals. B) **Correlation between cytokine PRS and time. using multiple thresholds reveals consistent trend**. Missing genotype rate ranged from 0.96, 0.9, 0.8, and 0.7. QTL P value for variants included in our PRS models ranged from 10^−3^, 10^−4^, 10^−5^, and 10^−6^. The color key indicates the range of −log10 P values of the Pearson correlation between PRS and time. Red and blue indicate positive and negative association, respectively.

Importantly, our results also show significant patterns in the changes of the production of specific cytokines during history. The resistance against intracellular pathogens increased after Neolithic with higher IFNγ responses (see Fig.1): indeed, it is known that Th1-IFNγ responses are crucial for the host defense against intracellular pathogens such as mycobacteria or *Coxiella* (Thakur et al., 2019). In addition, the resistance to the extracellular pathogens *C. albicans* and *S. aureus* is also increased after this Neolithic era, with TNFα and IFNγ production increasing steadily after. These two cytokines are very well known to be important for anti-*Candida* and anti-*Staphylococcus* host defense (Chan et al., 2018; Domínguez-Andrés et al., 2017). On the other hand, a different pattern emerges in relation with the IL1β-IL6-IL17 axis: the production of these cytokines is decreasing after Neolithic (see Figs. 1a and 1b). In this context, the decrease through time of poly I:C induction of cytokines, as a model of viral stimulation, is intriguing but potentially very important: many important viruses such as influenza and coronaviruses (SARS, MERS, and SARS-CoV-2) exert life-threatening effects through induction of cytokine-mediated hyperinflammation (also termed “cytokine storm’) (Tay et al., 2020): evolutionary processes to curtail this exaggerated responses are thus likely to be protective, and tolerance against viruses become a host defense mechanism (Diard and Hardt, 2017).

These evolutionary genetic adaptations to pathogens throughout human history greatly influence the way we respond to multiple diseases in modern times as well. To assess these effects, we calculated the PRS associated with the risk of several highly prevalent immune mediated diseases. The first focus was on common infectious diseases such as malaria, HIV-AIDS, tuberculosis and chronic viral hepatitis; we calculated the changes in susceptibility to these diseases in the last 50.000 years of human history, based on summary statistics from genome-wide association studies (GWAS) databases available from the literature (Fig. S3). Our results show that humans are becoming more resistant to these diseases, with the notable exception of tuberculosis, whose risk score remained stable along the period studied (Fig. 2). These results suggest that humans have built up a genetic makeup which made them more resistant to a variety of microbes. The pattern of this adaptation is very interesting as well, with a suggested decrease of susceptibility to malaria especially in the last 10.000 years. The reason for this accelerated resistance after Neolithic might be linked to higher disease prevalence due to increased populations density, as otherwise *Plasmodium* parasites are known to have circulated in Africa since at least the Paleogene 30 million years ago (Poinar, 2005), and we have likely inherited it from gorillas(Liu et al., 2010). Intriguingly, we also observe a strong decrease in susceptibility to HIV: this is a contemporary pathogen, therefore this signal could be due to common genetic and immune pathways with other infections that were present in human populations. The increased resistance to HIV in Europeans may be derived from selective pressures induced by other pathogens such as *Yersinia pestis* (Duncan et al., 2005). Our data suggest on the other hand that the source of this increased resistance is even older.

**Figure 2:**
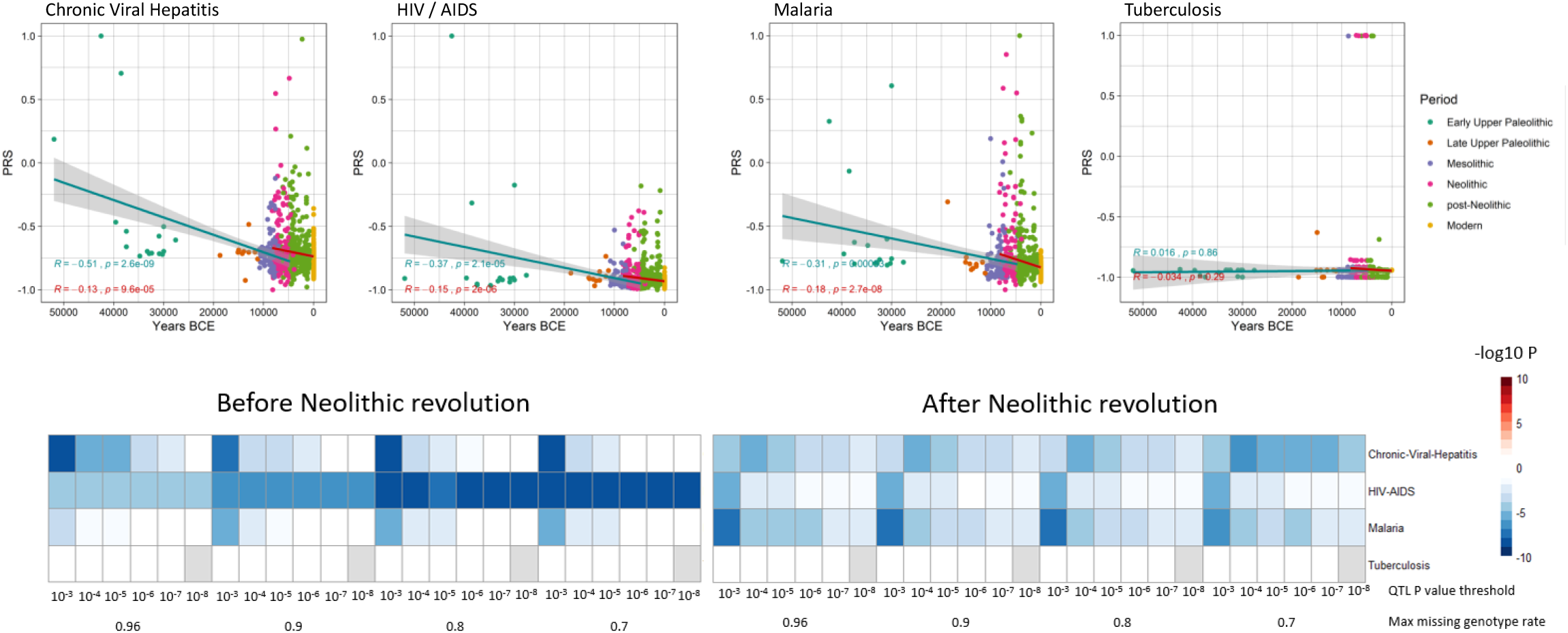
Infectious disease risk PRS scores decrease with time, except tuberculosis. **A)** Samples are colored by broad age period. The blue regression lines show PRS before the Neolithic revolution remained relatively constant for all traits whereas the red regression lines show that the correlation after the start of the Neolithic period changed significantly. The threshold of max missing genotype per sample was 0.96 and QTL P value cutoff was 10^−3^. MSUC: Monosodium urate crystals. **B) Correlation between disease PRS and time using multiple thresholds reveals consistent trend**. Missing genotype rate ranged from 0.96, 0.9, 0.8, and 0.7. QTL P value for variants included in our PRS models ranged from 10^−3^, 10^−4^, 10^−5^, 10^−6^, 10^−7^, and 10^−8^. The color key indicates the range of −log10 P values of the Pearson correlation between PRS and time. Red and blue indicate positive and negative association, respectively.

In contrast, the lack of genetic adaptation in the susceptibility to tuberculosis is intriguing. This surprising finding may be explained by a concept in which *M. tuberculosis* is at the same time a pathogen and a symbiont, in which latent infection enhances the resistance against other pathogens and this is why our immune system tolerates mycobacterial presence (Pai et al., 2016). In this regard, individuals with latent TB exhibit enhanced macrophage functions that may protect against other pathogens through the induction of trained immunity (Joosten et al., 2018). In this context humanity may not be adapting to tuberculosis because increased resistance against mycobacteria is not evolutionarily advantageous. All in all, these results suggest that the risk of suffering infectious diseases has steadily decreased at least for the last 50000 years as a result of the selection of genetic variants which confer resistance to infections.

It has been proposed that the increased prevalence of inflammatory and autoimmune diseases is associated with the immune-related alleles that have been positively selected through evolutionary processes to protect against infections, hence the contrasting differences in the prevalence of autoimmune diseases between populations results from diverse selective pressures (Ramos et al., 2015). In line with this, it has been hypothesized that genetic variants associated with protection against infectious agents are behind the increased prevalence of autoimmune diseases in populations with low pathogen exposure, such as Europeans (Fumagalli et al., 2011; Raj et al., 2013). To study the changing patterns of susceptibility to autoimmune and inflammatory disease during history, we used publicly available summary statistics from GWAS of digestive tract-related autoimmune and inflammatory diseases and arthritis-related diseases (Fig. S4) and calculated the PRS for each of samples under study. Interestingly, we observed a robust increase of the genetic variants related with the development of inflammatory diseases in the digestive tract after the Neolithic revolution (Fig. 3). PRS scores associated with celiac disease, Crohn’s disease, ulcerative colitis and inflammatory bowel disease, were strongly associated with the age of the samples, regardless of the P value thresholds or the missing genotype rates used for PRS calculation, showing the robustness of these results (Fig. S2). The fact that especially intestinal inflammatory pathology is increased after a historical event that fundamentally modified human diet is unlikely to be an accident. Our results are in line with earlier research demonstrating that variants in genes important for immune responses and involved in celiac disease pathophysiology (such as IL-12, IL-18RAP, SH2B3) are under strong positive selection (Zhernakova et al., 2010). The reasons for the selection pressure on these genes are not completely understood, but an advantage for host defense has been suggested (Zhernakova et al., 2010).

**Figure 3:**
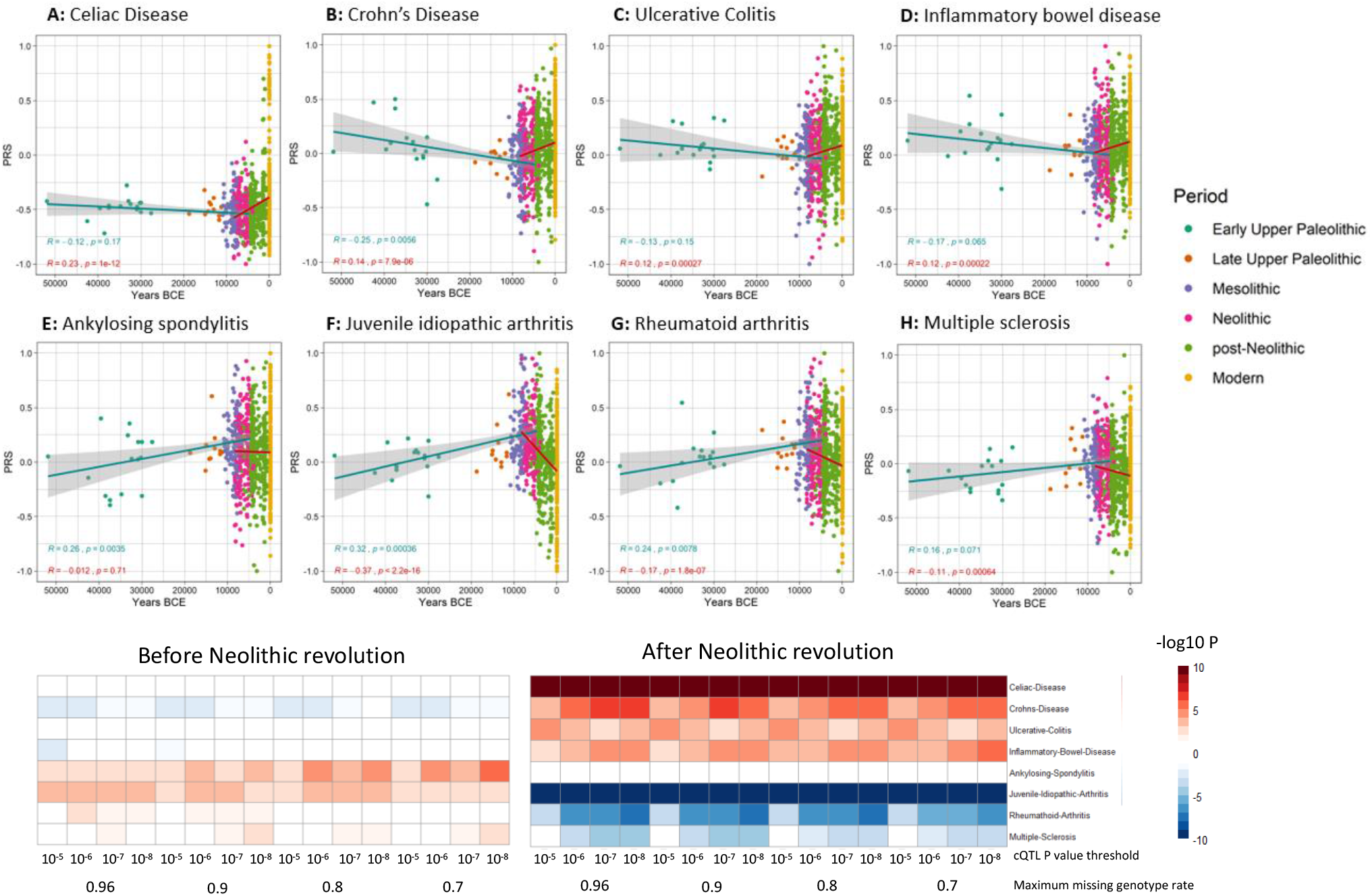
Correlation between auto-immune and inflammatory disease PRS and time. **A)** Samples are colored by broad age period. The blue regression lines show PRS before the Neolithic revolution remained relatively constant for all traits whereas the red regression lines show that the correlation after the start of the Neolithic period changed significantly. The threshold of max missing genotype per sample was 0.96 and QTL P value cutoff was 10^−4^. MSUC: Monosodium urate crystals. **B) Correlation between disease PRS and time using multiple thresholds reveals consistent trend**. Missing genotype rate ranged from 0.96, 0.9, 0.8, and 0.7. QTL P value for variants included in our PRS models ranged from 10^−5^, 10^−6^, 10^−7^, and 10^−8^. The color key indicates the range of −log10 P values of the Pearson correlation between PRS and time. Red and blue indicate positive and negative association, respectively.

In contrast to intestinal inflammation, the PRS of traits linked with juvenile-idiopathic arthritis, rheumatoid arthritis and multiple sclerosis shows a decrease in genetic susceptibility with the age of the sample after the Neolithic revolution. For pre-Neolithic periods, these patterns had little impact with decreasing PRS for digestive tract diseases and increasing PRS for ankylosing spondylitis and juvenile idiopathic arthritis. A strong decrease in susceptibility to juvenile idiopathic arthritis, rheumatoid arthritis and multiple sclerosis is seen after the Neolithic period (see Fig. 3). This is likely linked to the decreased production of the IL-1/IL-6/IL-17 axis described in Fig. 2, which is particularly important in the pathophysiology of these disorders (Akioka, 2019; Mei et al., 2011).

The significant changes in cytokine production and disease susceptibility in European populations after the Neolithic can be due to selective processes on the one hand (as described above), but also with important demographic changes due to migrations of human communities such as the Anatolians (in Neolithic) or the Yamnaya populations from the Pontic steppe (during the Bronze Age) (Racimo et al., 2020). In this regard, several loci associated with inflammatory disease displayed a group alleles linked with Crohn’s disease, celiac disease and ulcerative colitis in Neolithic Aegeans, the community who spread farming across Europe (Hofmanová et al., 2016), with several of these alleles showing signs of positive selection in modern Europeans (Raj et al., 2013). In addition, the gene expression PRS of several cytokines based on the *cis*- and *trans*-eQTLs from the eQTLGen Consortium (https://www.eqtlgen.org/) displayed a very strong association with time for TNFα after the Neolithic revolution (Fig. 4).

**Figure 4:**
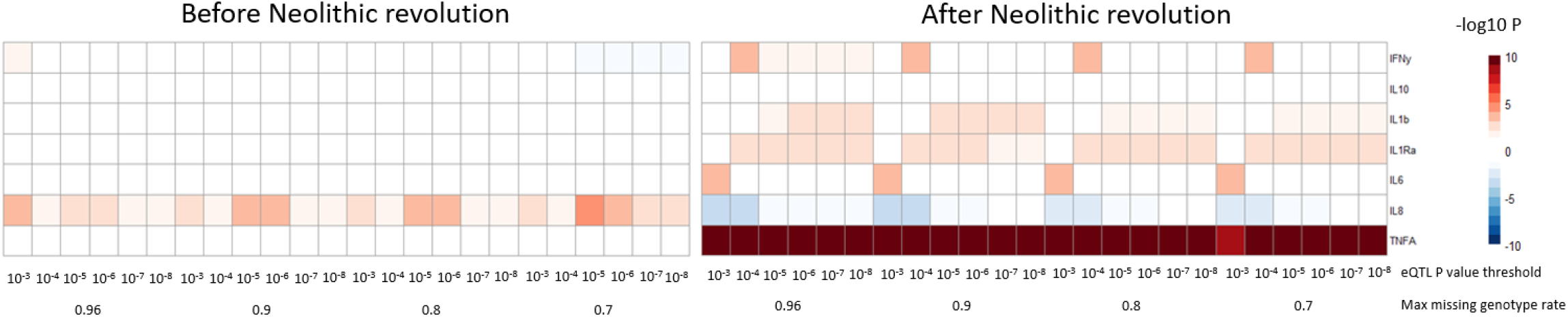
Cytokine gene expression PRS scores using cis- and trans-eQTLs correlated with time. Most notably is the highly significant increase in *TNFA* gene expression PRS over time following the Neolithic revolution. Prior to the Neolithic revolution an increase in *IL8* gene expression PRS can be observed which shifts to a decreasing trend after the Neolithic revolution. Both *IL1B* and *IL1RN* gene expression show a slight increase in PRS over time after the start of the Neolithic revolution. Missing genotype rate ranged from 0.96, 0.9, 0.8, and 0.7. QTL P value for variants included in our PRS models ranged from 10^−3^, 10^−4^, 10^−5^, 10^−6^, 10^−7^, and 10^−8^. The color key indicates the range of −log10 P values of the Pearson correlation between PRS and time. Red and blue indicate positive and negative association, respectively.

Collectively, our results show that the advent of the Neolithic era was a turning point for the evolution of immune-mediated traits in European populations, driving the expansion of alleles that favor the development of tolerance against intracellular pathogens and promote inflammatory responses against extracellular microbes. This is associated with a higher presence of genetic traits related with inflammatory and auto-immune diseases of the digestive tract and a lower number of alleles linked with the development of arthritis. Further research should compare the trends in different populations that have been exposed to different environments across the planet and clarify the influence of ancestry, time, rural vs. urban lifestyle to shed light on the influence of the infectious environment in genetics and human evolution.

## Acknowledgements

MGN was supported by an ERC Advanced Grant (833247) and a Spinoza Grant of the Netherlands Organization for Scientific Research. YL was supported by an ERC Starting Grant (948207) and the Radboud University Medical Centre Hypatia Grant (2018) for Scientific Research. JB was supported by PID2019-110933GB-I00 (AEI/FEDER, UE) MINECO, Spain.

## Author contributions

JDA, YK, OBB and MJ designed and performed experiments and analysed the data. JDA and YK wrote the first draft of the manuscript with all authors contributing to writing and providing feedback. MJ, CJX, JWMvdM, LABJ and JB provided guidance and advice. YL and MGN conceived ideas, designed experiments, offered supervision and oversaw the research program.

## Methods

### Cohort selection

Ancient DNA genotype data was downloaded from version 37.2 of the published aDNA genotype database, compiled by and available on the David Reich Lab website (https://reich.hms.harvard.edu/downloadable-genotypes-present-day-and-ancient-dna-data-compiled-published-papers). The ancient DNA samples consisted of pseudo-haploid genotype data. This was due to the low genotyping coverage. Samples with variant missingness above 96 percent were filtered out using Plink (Purcell et al., 2007). This was done in order to remove outliers with extremely low coverage. Only samples within Europe were used for this study, these samples were selected based on their geographic location, that is latitude (within 35 and 70 degrees north) and longitude (within 10 degrees west and 40 degrees east). Samples without a carbon-dated age were also filtered out. We also selected 250 European samples from the 1000 genomes project phase 3. Only variants present in both the ancient samples and the modern samples were retained. This resulted in a dataset of 827 ancient samples and 250 modern samples containing 1233013 variants.

### Carbon-dated sample origin and geographical location

Both carbon-dated age of origin as well as latitudinal and longitudinal data was available for these 827 ancient European samples. Broad time periods were assigned to these samples with the Early Upper Paleolithic era for all samples originating from before 25000 years before the common era standardized to 1950 (BCE). The Late Upper Paleolithic era follows until 11000 BCE. The Mesolithic era ranges from 11000 to 5500 BCE. The Neolithic era ranges from 8500 to 3900 BCE, and the Post-Neolithic era ranges from 5000 BCE and more recent ages. Using the geographical data in combination with archeological clues and the genetic data, the broad time period of origin was also available for samples that were dated to a point in time with overlapping broad time periods. This allowed the samples to be classified as either Early Upper Paleolithic, Late Upper Paleolithic, Mesolithic, Neolithic, or Post-Neolithic. The sample age of the 250 modern European samples was set to 0.

### Summary statistics of GWAS and cytokine QTLs

Summary statistics for complex traits were obtained from the UK Biobank (Bycroft et al., 2018) and the GWAS catalog (MacArthur et al., 2017) last accessed on 29^th^ of March 2020. The stimulated cytokine response summary statistics from the 500FG cohort of the HFGP were used (Li et al., 2016). Some complex traits had multiple different sets of summary statistics available. In these cases, the data which was more recent and used bigger cohorts that were either of European or mixed (European and Asian) ancestry were selected. The variants of these summary statistics were then filtered by only keeping bi-allelic variants. Most aDNA genotypes available are pseudo-haploid as a consequence of their lower sample quality. We excluded ambiguous SNPs (A/T and C/G) in order to prevent errors due to strand flips present in these pseudo-haploid samples.

### Polygenic Risk Scores (PRS) calculation

Polygenic risk scores were then calculated by first intersecting the filtered variants from the summary statistics with the variants present in the DNA samples. Starting at the most significant variant, all variants within a 250kb window around that variant were excluded until no variants remained. We then multiplied the dosage of these variants with the effect size and these values were summed. If a variant is missing in a sample the dosage is substituted with the average genotyped dosage for that variant within the entire dataset. This way the PRS is not skewed in any specific direction. The formula for this is described below with the score *S* being the weighted sum of a variant’s dosage *X*_*n*_ multiplied by its associated weight or beta *β*_*n*_ calculated using *m* variants.

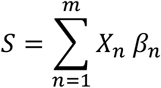

### Piecewise correlation analysis

We constructed piecewise linear models for each trait by separating the samples into two groups. These two groups consisted of all samples preceding the Neolithic era and those of the Neolithic era and later respectively. We correlated PRS with the carbon dated age of our samples. We then multiplied the −log10 of the correlation P values with the sign of the correlation coefficients.

### Robustness of results

In order to test the robustness of our results we calculated PRS using multiple different P value thresholds for QTL inclusion. We used P value thresholds from 10^−3^ to 10^−8^ for the complex traits obtained through GWAS catalog and the UK Biobank. The thresholds used for the stimulated cytokine responses ranged from 10^−3^ to 10^−6^. We also calculated PRS using different variant missingness thresholds. This means we removed samples with a variant missingness rate higher than 96, 90, 80, or 70 percent. All of the results from the piecewise linear models were then used to create a heatmap depicting the consistency and robustness of our observed correlations.

Additionally, various window-sizes were used for clumping the QTL’s and LD based clumping was also performed excluding variants with an LD greater than 0.2 compared to our lead SNP within a window. In order to see whether our observations were due to sample imbalances between the pre-Neolithic period and the later periods samples originating from the Neolithic period and later were randomly down-sampled to the same number of samples as the pre-Neolithic samples. Correlation coefficients between PRS and sample age were then recalculated for the Neolithic and younger samples and compared to the coefficients obtained using all Neolithic and younger samples.

## Figures

**Figure S1:**
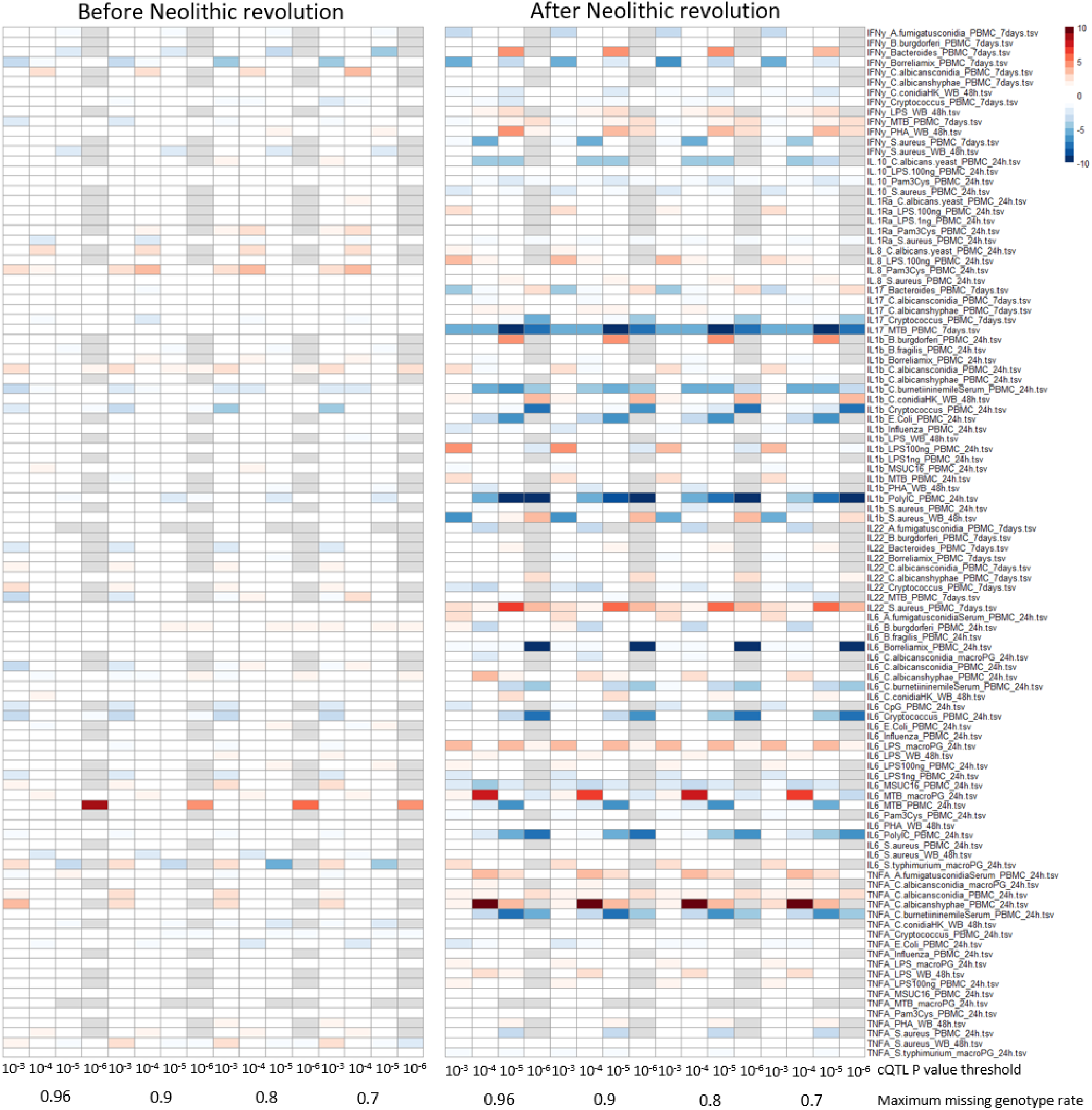
Correlation between cytokine PRS and time. Missing genotype rate ranged from 0.96, 0.9, 0.8, and 0.7. QTL P value for variants included in our PRS models ranged from 10^−3^, 10^−4^, 10^−5^, and 10^−6^. The color key indicates the range of − log10 P values of the Pearson correlation between PRS and time. Red and blue indicate positive and negative association, respectively.

**Figure S2:**
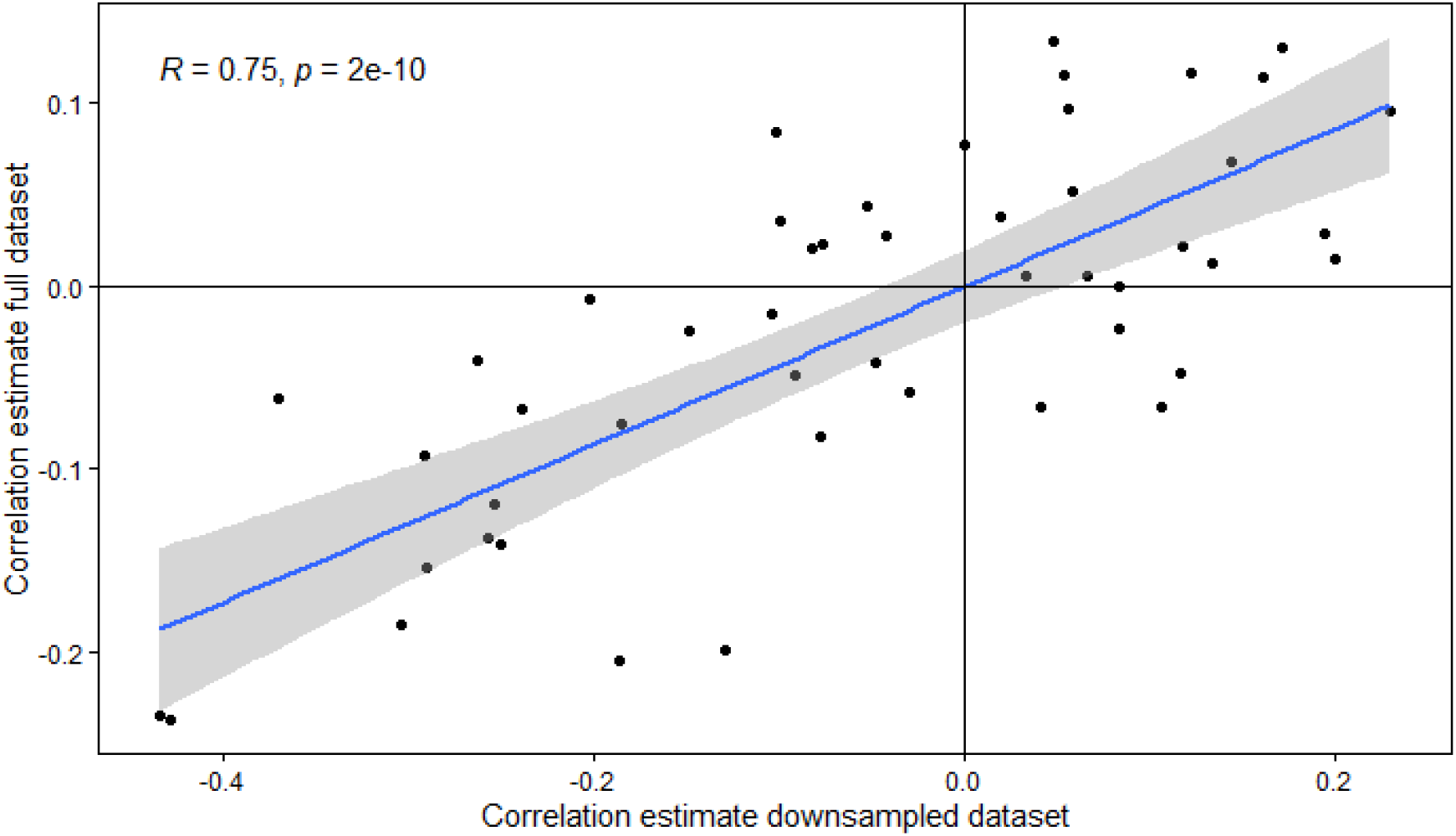
Robustness of correlation coefficients post Neolithic independent of sample size. Changes in PRS following the Neolithic revolution remain consistent after down-sampling samples from after the start of the Neolithic period to the same amount as samples before the Neolithic period. The lower number of samples reduces the power which reduces the amount of significant correlations but does not influence the direction of changes in PRS which were previously identified as significant.

**Figure S3:**
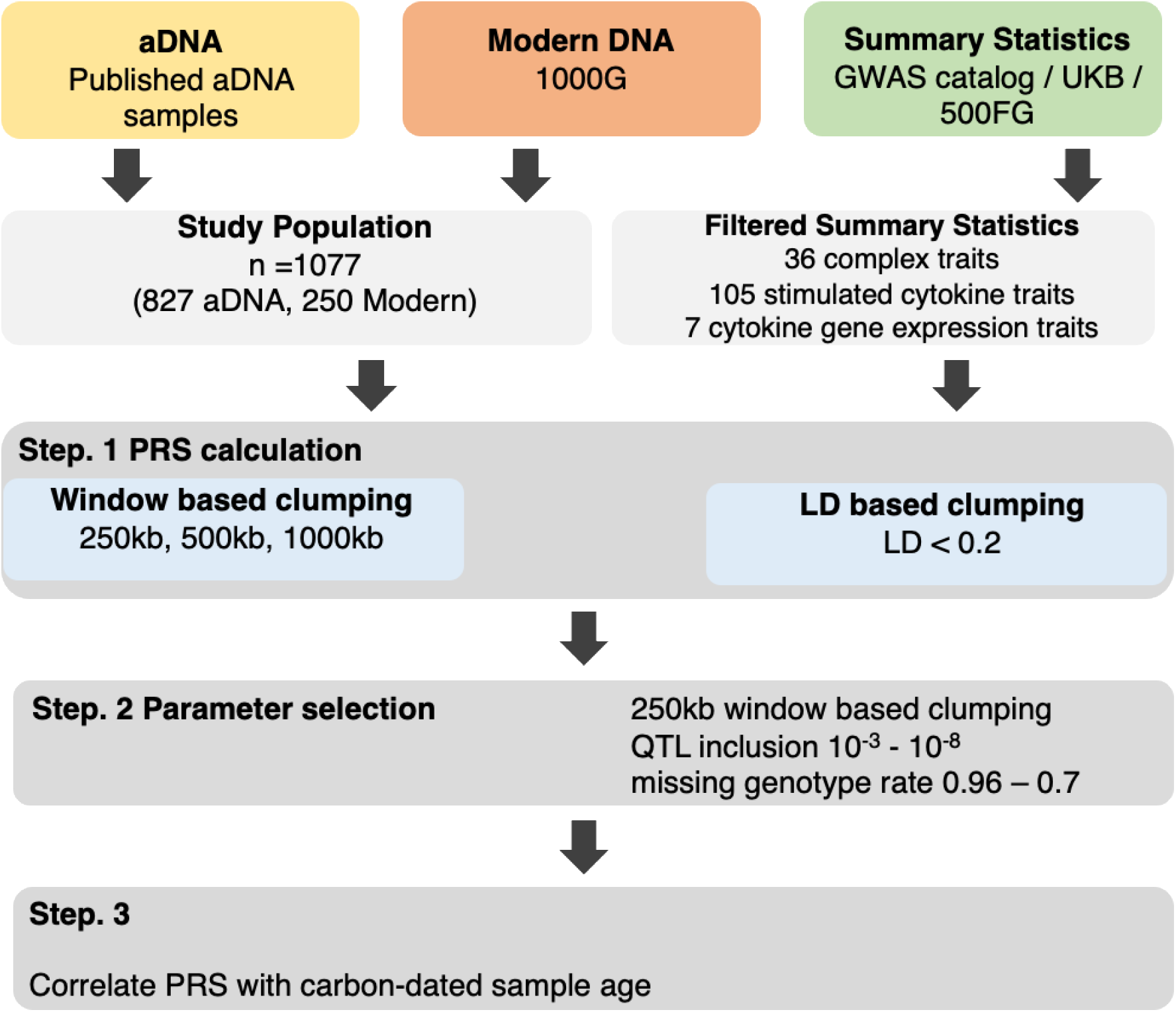
Both aDNA and modern DNA samples of European individuals were used in combination with summary statistics from predominantly European populations to calculate PRS of immune-related traits. This was done at various threshold combinations before correlating the scores with the sample age.

**Figure S4:**
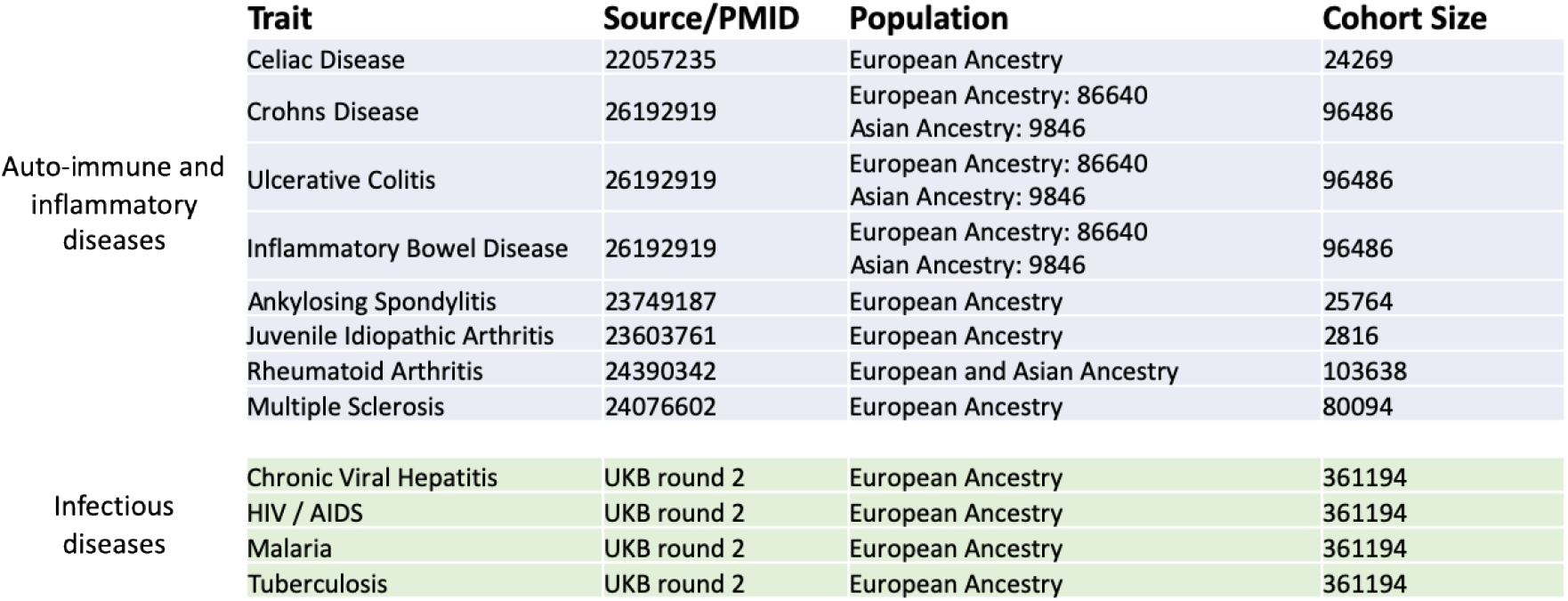
GWAS summary statistics and cohorts used for PRS calculation. Traits were separated into two categories: Auto-immune and inflammatory diseases-related traits, and infectious diseases-related trait. GWAS summary statistics from predominantly European populations were selected.

**Table 1:**
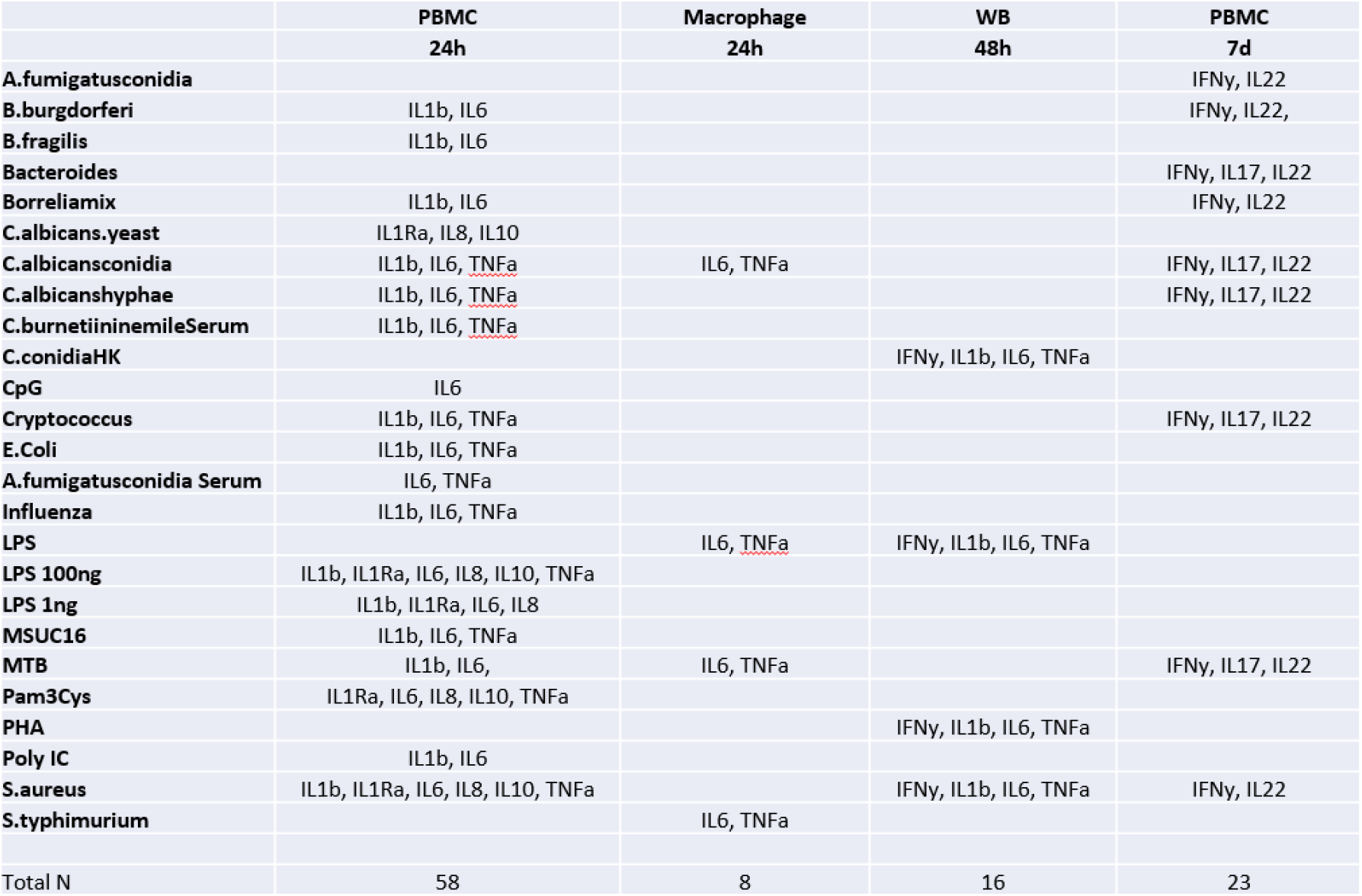
Overview of the stimulus, cytokine, and timepoint combinations. In total105 unique stimulated cytokine traits were available using various types of stimuli measuring both the innate and adaptive immune response.

